# Distinct modes of coupling between VCP, an essential unfoldase, and deubiquitinases

**DOI:** 10.1101/2024.09.08.611915

**Authors:** Lauren E. Vostal, Noa E. Dahan, Wenzhu Zhang, Matthew J. Reynolds, Brian T. Chait, Tarun M. Kapoor

## Abstract

Errors in proteostasis, which requires regulated degradation and recycling of diverse proteins, are linked to aging, cancer and neurodegenerative disease (1). In particular, recycling proteins from multiprotein complexes, organelles and membranes is initiated by ubiquitylation, extraction and unfolding by the essential mechanoenzyme VCP (2–4), and ubiquitin removal by deubiquitinases (DUBs), a class of ∼100 ubiquitin-specific proteases in humans (5, 6). As VCP’s substrate recognition requires ubiquitylation, the removal of ubiquitins from substrates for recycling must follow extraction and unfolding. How the activities of VCP and different DUBs are coordinated for protein recycling or other fates is unclear. Here, we employ a photochemistry-based approach to profile proteome-wide domain-specific VCP interactions in living cells (7). We identify DUBs that bind near the entry, exit, or both sites of VCP’s central pore, the channel for ATP-dependent substrate translocation (8–10). From this set of DUBs, we focus on VCPIP1, required for organelle assembly and DNA repair (11–13), that our chemical proteomics workflow indicates binds the central pore’s entry and exit sites. We determine a ∼3Å cryo-EM structure of the VCP-VCPIP1 complex and find up to 3 VCPIP1 protomers interact with the VCP hexamer. VCPIP1’s UBX-L domain binds VCP’s N-domain in a ‘down’ conformation, linked to VCP’s ADP-bound state (2, 14), and the deubiquitinase domain is positioned at the central pore’s exit site, poised to remove ubiquitin following substrate unfolding. We find that VCP stimulates VCPIP1’s DUB activity and use mutagenesis and single-molecule mass photometry assays to test the structural model. Together, our data suggest that DUBs bind VCP at distinct sites and reveal how the two enzyme activities can be coordinated to achieve specific downstream outcomes for ubiquitylated proteins.

## Introduction

Processing of ubiquitylated proteins for degradation and recycling depends on VCP (valosin-containing protein, or p97), an AAA (ATPases associated with diverse cellular activities) unfoldase (2–4). VCP extracts polyubiquitylated proteins from macromolecular complexes, organelles, and membranes and unfolds them, ATP-dependent processes needed for the degradation or recycling of proteins (2–4). Consistent with these essential functions, VCP overexpression or mutations are linked to neurodegenerative disorders, such as multisystem proteopathies (15) and vacuolar tauopathy (16), and cancer (17), and chemical inhibitors, as well as activators, of VCP have been developed as potential therapeutics (17–19). Structurally, VCP consists of a globular N-terminal domain (N), two ATPase domains (D1/D2), and a disordered C-terminal tail (14). The N- and C-terminal domains regulate enzymatic activity through cofactor binding (20) or phosphorylation (21–23) and can contribute to chemical activation (18). In particular, the N domain adopts ‘up’ or ‘down’ conformations, coaxial or coplanar with the D1 domains, respectively, depending on VCP’s nucleotide state (2, 14). Extraction and unfolding occurs through recognition of K48-linked ubiquitylated substrates by cofactors and subsequent ATP-dependent threading of substrate and ubiquitin through a pore at the center of the ring-shaped VCP hexamer (hereafter, central pore) (8–10). Unfolded polyubiquitylated substrates are released from VCP and targeted to the proteasome (8, 24, 25). For recycling, the ubiquitin chain must be removed after substrate recognition, extraction and unfolding by VCP. However, we do not know how VCP activity is coupled to the removal of ubiquitin from substrates.

DUBs are a class of proteases, separated into seven families according to their catalytic domains, that recognize and selectively cleave ubiquitin. Ubiquitin can assemble into covalent polymeric chains through internal lysine residues, resulting in distinct structures (e.g., branched or linear) that regulate diverse cellular pathways, including proteostasis, cell signaling, and cell division (1, 26). Consistent with the critical roles of ubiquitylation, DUBs have been shown to modulate the levels of oncogenes or tumor suppressor genes and regulate cellular pathways including the ubiquitin-proteasome system (UPS) and mitophagy, and are currently attractive targets for the development of cancer and neurodegenerative disease therapies (6, 27). A subset of these DUBs have been shown to interact with VCP in biochemical experiments (using recombinant proteins or overexpression of DUBs in human cell lysate), most notably ataxin-3, YOD1, USP13, and VCPIP1 (28–30). The only structural data supporting a direct VCP-DUB interaction is a crystal structure of minimal truncated constructs of the *S. cerevisiae* DUB OTU1 (YOD1 in humans) UBX-L domain, a noncatalytic VCP-interacting motif, and the VCP N domain alone (31). Based on these data, current models suggest DUBs, which recognize a folded ubiquitin, bind the near the entry to the substrate unfolding site in VCP and can cleave a subset, but not all the ubiquitins linked to substrates (8). However, we lack structural models for how the active site of any DUB is coupled to VCP’s unfolding and how ubiquitins can be removed after substrates have been unfolded.

Here, we employ a chemical proteomics workflow combining amber suppression, photo-crosslinking, and quantitative mass spectrometry to profile direct, domain-specific VCP interactions in living human cells. We identify DUBs that bind near the entry, exit, or both sites of VCP’s central pore. We determine a cryo-electron microscopy (cryo-EM) structure of the VCP-VCPIP1 complex that reveals how VCPIP1 interacts with both sites of VCP. We find that the VCPIP1 deubiquitinase domain is positioned below the exit of the VCP central pore to remove ubiquitin from unfolded or extracted substrates. We test this model using mutagenesis and mass photometry and show that VCPIP1’s DUB activity is stimulated in the presence of VCP. Together, our findings suggest a model for how the coupling of VCP with DUBs can regulate distinct outcomes for polyubiquitylated substrates.

## Results

### A chemical proteomics approach identifies deubiquitinases that directly engage VCP in distinct modes

To analyze direct and domain-specific interactions between VCP and its associated proteins in living human cells, we adapted a chemical proteomics approach combining amber suppression, photo-crosslinking and quantitative mass spectrometry (7) (Fig. 1a). Briefly, amber suppression is used to incorporate 3’azibutyl-*N*-carbamoyl-lysine (AbK), a photo-crosslinkable amino acid, or N^ε^ -Boc -lysine (BocK) as a control, at a selected position in the protein of interest in HEK293T cells (Fig. 1a). Exposure of cells to UV light triggers diazirine-dependent crosslinking and covalently-linked complexes are isolated, using a two-step tandem affinity protocol with one step under denaturing conditions to minimize capture of non-covalent interactors. The use of C-terminal affinity tags enables purification of only full-length VCP that has incorporated the non-natural amino acid. Purified samples are then processed for mass spectrometry (Fig. 1a).

**Fig. 1:**
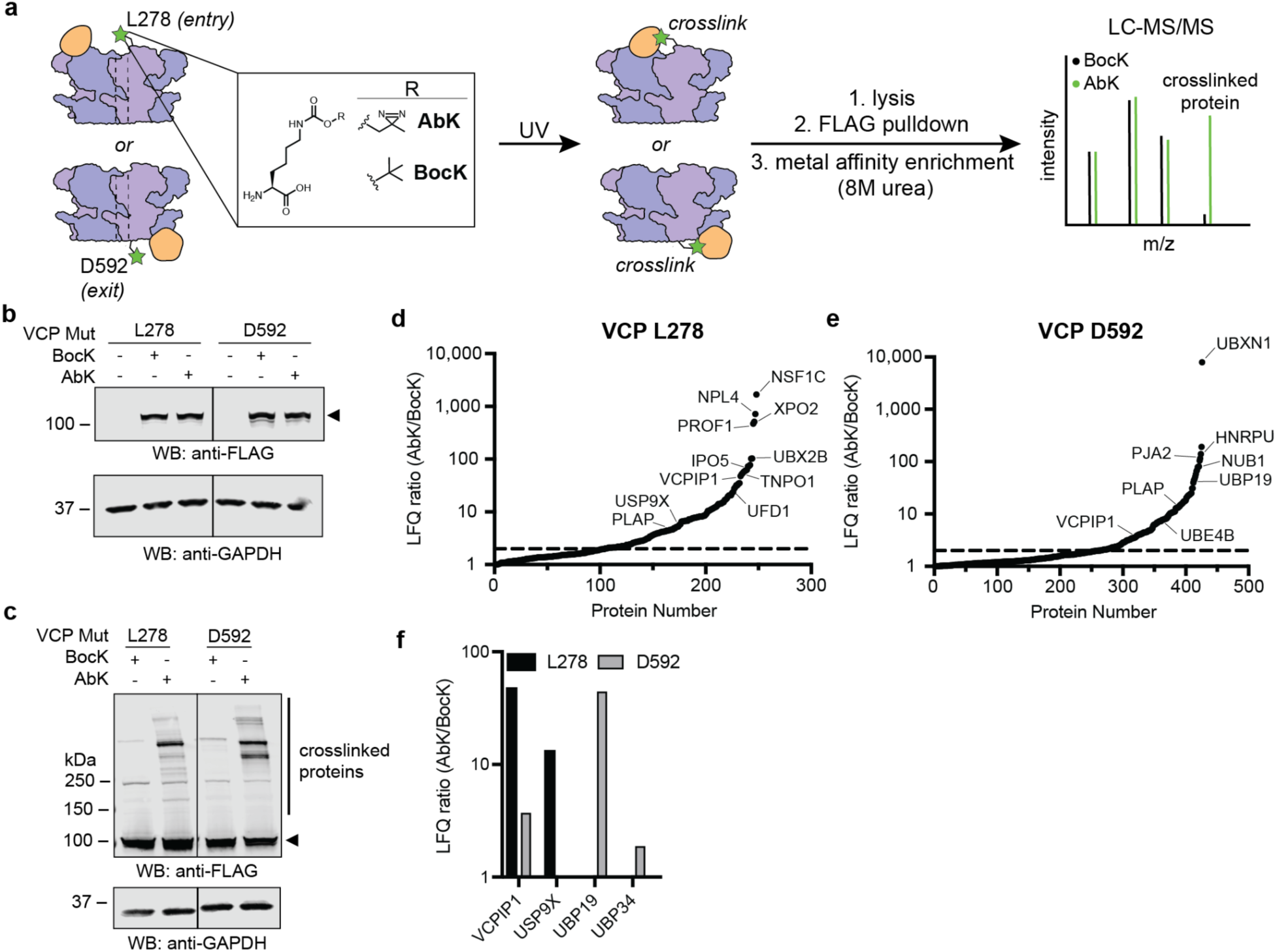
A chemical proteomics approach identifies deubiquitinases that engage VCP at the entry, exit, or both sites of its central pore. **a**, Schematic for the chemical proteomics approach. VCP (purple) is expressed in HEK293T cells with non-natural amino acids (AbK or BocK, green star) site-specifically incorporated near the central pore (dashed lines) by amber suppression. UV exposure initiates photo-crosslinking with associated proteins (orange), only in the presence of AbK. BocK serves as a control. Covalent complexes are purified using tandem-IP, with one step under denaturing conditions, and analyzed by LC-MS/MS. **b**, Western blots of HEK293T cells expressing VCP with non-natural amino acid (AbK or BocK) incorporated at L278 (VCP-L278-AbK or VCP-L278-BocK) or D592 (VCP-D592-AbK or VCP-D592-BocK) (arrowhead). **c**, Western blots of HEK293T cells expressing VCP-L278 or VCP-D592 in the presence of AbK or BocK (arrowhead) after exposure to UV light. d, Plot of fold change in label-free quantification for VCP-L278-AbK versus VCP-L278-BocK. Shown are proteins with fold change > 1 (dashed line = fold change > 2). **e**, Plot of fold change in label-free quantification for VCP-D592AbK versus VCP-D592BocK. Shown are proteins with fold change > 1 (dashed line = fold change > 2). f,Plot of fold change in label-free quantification for deubiquitinases identified in VCP-L278 (black) and VCP-D592 (grey) datasets. Shown are deubiquitinases with fold change > 2 (4 of 8 total) in **d** and **e**.

To profile DUBs that interact with VCP at the entrance or the exit to its central pore, we incorporated the non-natural amino acids at two sites. First, we selected residue L278, which is positioned in a loop near the entrance to the VCP central pore. In yeast VCP (Cdc48) the residue at the equivalent position, M288, is positioned at the interface between VCP and UN (PDB: 6OA9) (32). Second, we selected residue D592, which is positioned in a loop at the exit of the central pore. The non-natural amino acid p-benzoylphenylalanine (Bpa) has been incorporated at these sites in both VCP (33, 34) and Cdc48 (8, 32) in bacteria and yeast respectively, but not in human cells or for proteomics.

Western blots indicated full length VCP expression only in the presence of either AbK or BocK (hereafter, VCP-L278AbK, VCP-L278BocK, VCP-D592AbK, VCP-D592BocK) (Fig. 1b). Further, AbK- or BocK-modified VCP were expressed at levels lower than endogenous VCP (Supplementary Data Fig. 1a,b). Upon processing for photocrosslinking in the presence of AbK, VCP and higher molecular weight bands are observed, consistent with covalent protein adduction (Fig. 1c). Substantially fewer adducts were detected in the presence of BocK (Fig. 1c). Gratifyingly, the crosslinking pattern observed in VCP-D592AbK is different from that of VCP-L278AbK, suggesting covalent linkage to distinct interactors depends on the position AbK is incorporated in VCP (Fig. 1c).

We next performed liquid chromatography-tandem mass spectrometry (LC-MS/MS) on samples after photo-crosslinking and tandem-IP. We employed label-quantification (LFQ) and identified proteins enriched two-fold or greater in VCP-L278/D592AbK compared to VCP-L278/D592BocK. Importantly, enrichment levels of hits are not dependent on their cellular abundance (35) or molecular weight (Supplementary Data Fig. 1c,d).

We identified 144 proteins as hits that likely bind near the entry to the VCP central pore (Fig. 1d and Supplementary Table 1). Notably, NSF1C/p47, NPL4, UBX2B/p37, VCPIP1, and UFD1 are enriched (fold change(AbK/BocK) = p47: 1681; Npl4: 717; p37: 102; VCPIP1: 48; Ufd1: 24) (Fig. 1d and Supplementary Table 1), which is consistent with these cofactors interacting near the central pore entry site through a VCP-interacting domain or motif (e.g., UBX, UBX-L, or SHP) (36). Two DUBs are enriched at the entry of VCP’s central pore (fold change(AbK/BocK) = VCPIP1: 48; USP9X: 13) (Fig. 1d,f and Supplementary Table 1). We note that known VCP-binding DUBs, such as ataxin-3, YOD1, USP13, are not enriched in our datasets, suggesting that these DUBs may not bind VCP at the selected sites or require different cellular conditions (i.e., stress) for interaction to occur.

We observe 165 proteins that are enriched two-fold or greater in VCP-D592AbK compared to VCP-D592BocK (Fig. 1e and Supplementary Table 2). In this dataset we also observe enrichment of known VCP cofactors, including UBXN1, PLAP/PLAA, UBE4B, and VCPIP1 (fold change(AbK/BocK) = UBXN1: 7888; PLAP/PLAA: 15; UBE4B: 8; VCPIP1: 4) (Fig. 1e and Supplementary Table 2). PLAP/PLAA contains a PUL (PLAA, Ufd3p, and Lub1p) domain that recognizes the C-terminal HbYX motif in VCP(36). We identified three DUBs that bind VCP at the exit to its central pore (fold change(AbK/BocK) = UBP19: 44; VCPIP1: 4; UBP34: 2) (Fig. 1e,f and Supplementary Table 2). Together, our chemical proteomics workflow identifies a subset of DUBs that in living human cells interact with VCP’s central pore proximal to the entry, exit, or one that binds both sites.

### Structural analysis of the VCP-VCPIP1 complex reveals up to three VCPIP1 protomers binding the exit of VCP’s central pore

To characterize the VCP-VCPIP1 interaction we expressed and purified full-length human VCP and VCPIP1 using published procedures with modifications (11, 18, 21) (see Methods) (Fig. 2a, Supplementary Data Fig. 2a). VCPIP1 contains an OTU catalytic domain and a UBX-L domain (27, 36) (Fig. 2a). During each protein purification we removed any affinity tags used. Proteins eluted as monodisperse peaks by size exclusion chromatography with sizes corresponding to VCP hexamer and VCPIP1 monomer (Supplementary Data Fig. 2b).

**Fig. 2:**
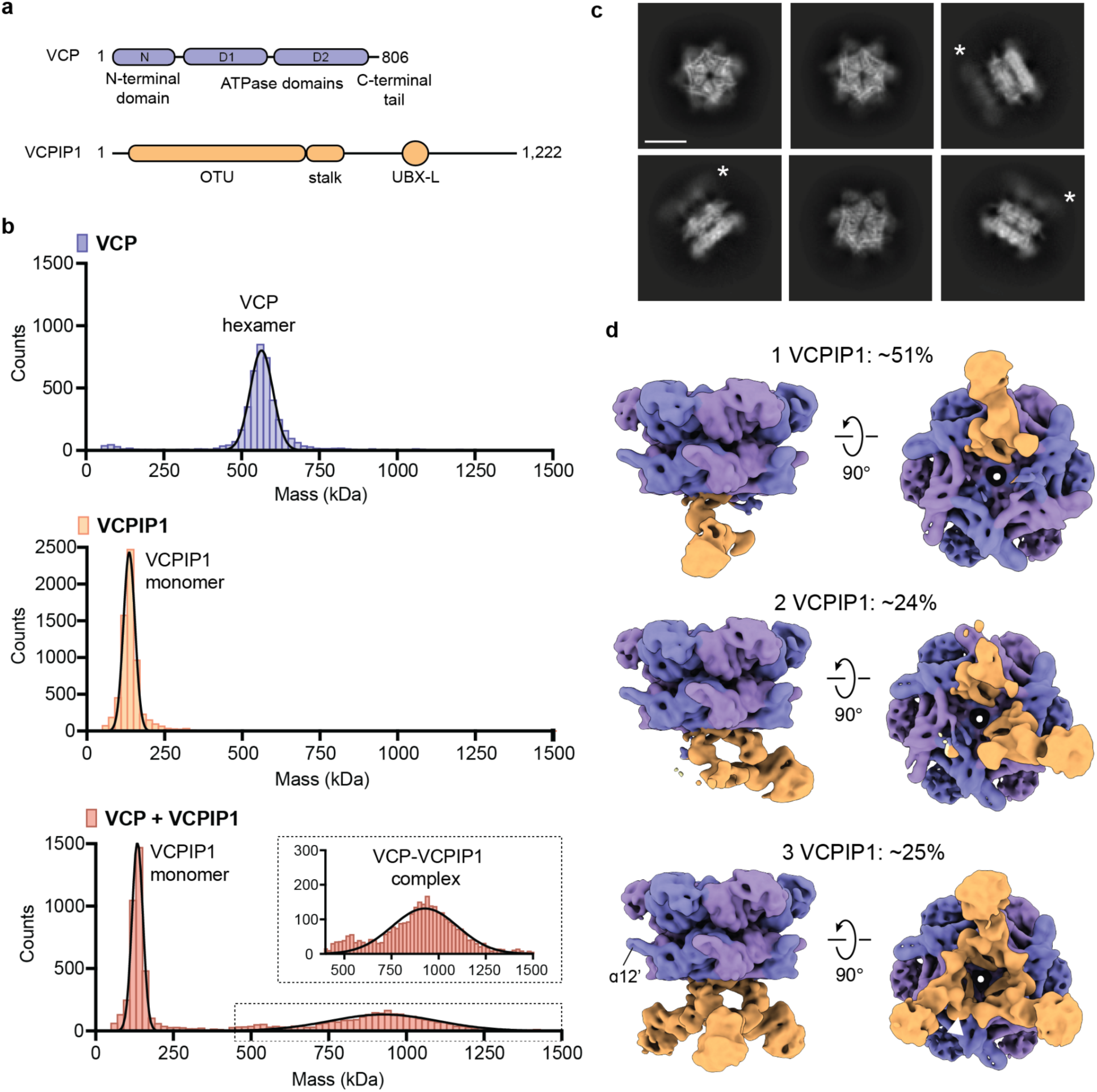
Cryo-EM analysis of the VCP-VCPIP1 complex reveals a binding interface at the exit of the VCP central pore. **a**, Domain maps of VCP (purple) and VCPIP1 (orange). **b**, Mass photometry of VCP-VCPIP1 complexes. Conditions include VCP WT alone (purple), VCPIP1 WT alone (orange), or VCP WT + VCPIP1 WT (red). The inset (dotted line) is a zoom of VCP WT + VCPIP1 WT between masses of 400 and 1500 kDa. **c**, 2D class averages following initial classification of the VCP-VCPIP1 dataset, showing the VCP hexamer and additional density for VCPIP1 (asterisk). Scale bar = 100 Å. **d**, Side and bottom (from below the VCP D2 domain) views of 3D classes with 1-3 VCPIP1 protomers (orange) bound to the VCP hexamer (blue and purple). Percentage of VCPIP1-bound particles (∼79% of total particles) in each class is shown. A class with no density for VCPIP1 is not shown (∼21% of total particles). *Bottom*, helix ɑ12’ is denoted and arrowhead (white) indicates density connecting VCPIP1 protomers.

We next used mass photometry (MP), which enables the determination of the mass of single molecules in solution(37). Using nanomolar concentrations of protein, we observe a gaussian distribution of masses for VCP (mean ± SD: 566 ± 59 kDa), consistent with a stable hexamer (Fig. 2b). VCPIP1 alone also revealed a gaussian distribution of masses (mean ± SD: 137 ± 52 kDa), corresponding to a monomer in solution. (Fig. 2b). When VCP hexamer was mixed with a four-fold excess of VCPIP1 monomer, we observe peaks for VCPIP1 monomer and VCP-VCPIP1 complex (Fig. 2b). Very little free VCP was detected. The broad peak representing the VCP-VCPIP1 complex was fit to a gaussian curve (mean ± SD: 930 ± 167 kDa) and corresponds to a stoichiometry of 3 VCPIP1 monomers bound to a single VCP hexamer. However, the standard deviation of this peak is on the order of 1 VCPIP1, which suggests that there is a mixture of stoichiometries under these conditions.

To examine the VCP-VCPIP1 complex, we used single-particle cryo-EM. VCP was mixed with excess VCPIP1 in the absence of nucleotide and samples were processed for cryo-EM, where micrographs revealed particles with various orientations (Supplementary Data Fig. 2c). Autopicked particles (∼2.9 million) were subjected to 2D classification, which yielded class averages that revealed stacked hexagons, as would be expected for VCP (Fig. 2c). We also observe an additional density associated with many of the stacked hexagons (Fig. 2c, asterisk).

We next separated particles into classes with or without the density associated with the stacked hexagons (Fig. 2d and Supplementary Data Fig. 3a). At this stage we were able to rigid body fit a model for VCP (PDB: 8OOI) (38) into the density. The assignment of VCP’s D2 ring was supported by helix ɑ12’, a prominent feature of the small subunit of the D2 ATPase domain (20, 39) (Fig. 2d, asterisk and Supplementary Data Fig. 3b). Our density agrees with the docked model at the N-domains, which are in their ‘down’ position, consistent with a state in which ADP is bound in the D1 domain (40) (Supplementary Data Fig. 3b).

With these assignments, we establish VCP as a homohexamer with stacked D1 and D2 domains, as has been previously reported (2, 14) (Fig. 2d). The additional density below the VCP D2 domain was too large and globular to be assigned to VCP’s C-terminal region, which is not completely accounted for in earlier structures (38, 40, 41). We assigned this density to VCPIP1. 3D classes with 1-3 VCPIP1 protomers comprise ∼79% of cleaned particles (∼40, 19, and 20%, respectively), consistent with results from MP experiments (Fig. 2b,d). In the class with three VCPIP1 protomers, we observe density connecting the protomers (Fig. 2d, arrowhead). In all classes, we observe one VCPIP1 protomer at the interface between two adjacent VCP protomers (Fig. 2d). Together, our cryo-EM data reveal a VCP-VCPIP1 structure in which up to three globular domains of the VCPIP1 protomer are positioned near the exit to VCP’s central pore.

### The VCPIP1 UBX-L domain binds the VCP N-domain and contributes to complex formation

We next identified an asymmetric unit containing one VCPIP1 and two VCP protomers (Figs. 2d and 3a). We next employed our second strategy including C3 symmetry expansion and masked 3D classification around the asymmetric unit (Supplementary Data Fig. 3a). This generated an overall map of ∼3.0 Å resolution (Fig. 3a and Supplementary Data Fig. 4a,b). In high resolution regions we observe density corresponding to secondary structure elements and side chains (Supplementary Data Fig. 4c). A subset of the map had high noise regions, which we low pass filtered to 7-8 Å resolution to generate a composite map (Supplementary Data Fig. 4d) (also see below).

**Fig. 3:**
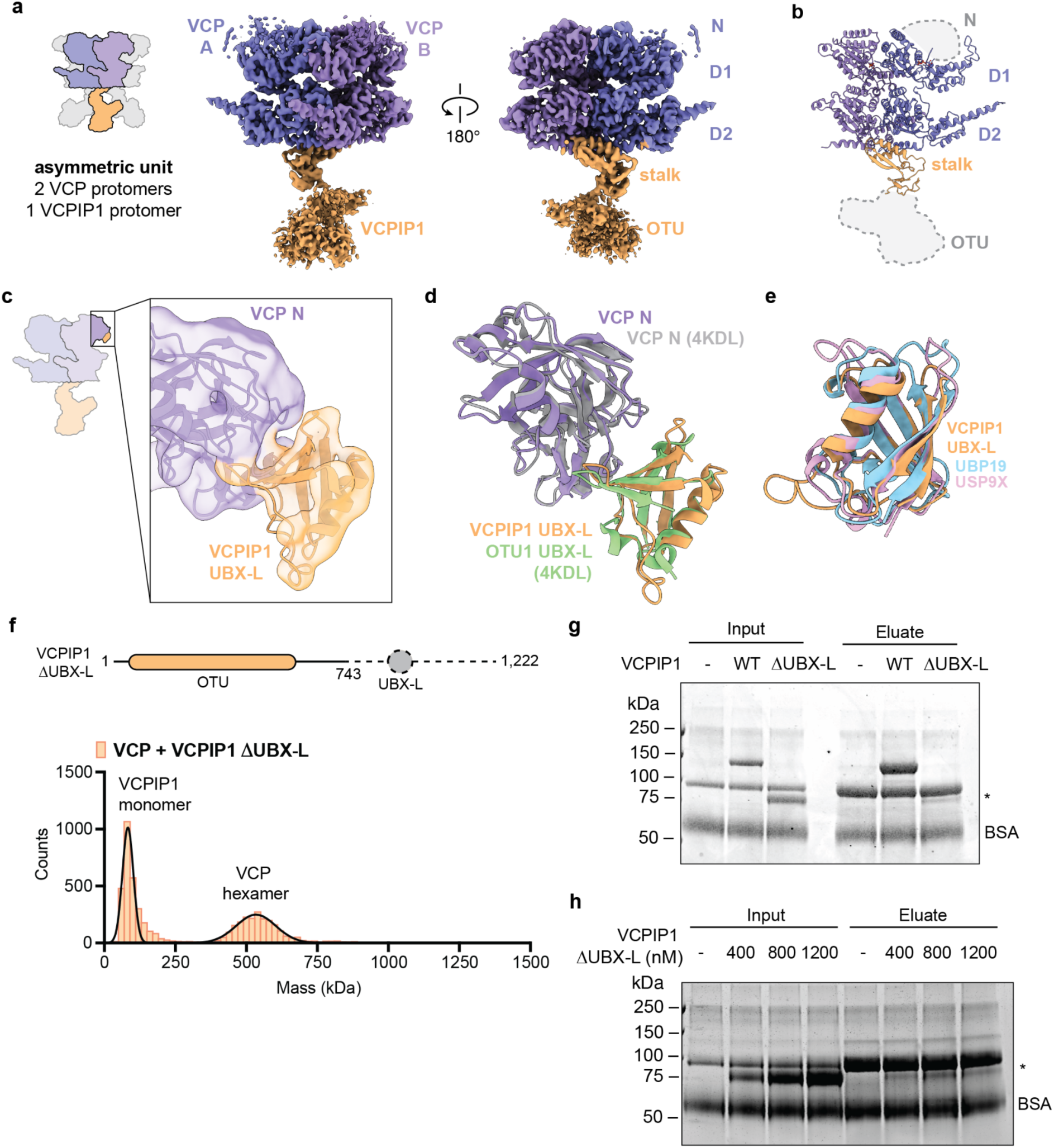
A second interaction with the regulatory VCP N-domain is sufficient for VCP-VCPIP1 complex formation. **a**, *Left,* Schematic for the asymmetric unit obtained after C3 symmetry expansion. The asymmetric unit consists of two VCP protomers (blue and purple) and one VCPIP1 protomer (orange). *Right*, Cryo-EM map of the VCP (blue and purple)-VCPIP1 (orange) complex in two different views. **b**, Model generated from the cryo-EM map of the VCP-VCPIP1 complex. The model consists of VCP D1 and D2 domains (blue and purple) and the VCPIP1 “stalk” (orange). The VCP N and VCPIP1 OTU domains (shaded grey with dashed lines) were not modeled at this stage due to flexibility (see below). **c**, *Left*, Schematic for the VCP (blue and purple)-VCPIP1 (orange) asymmetric unit with a box around the VCP N-VCPIP1 UBX-L interaction. *Right*, Cryo-EM map and model of the VCP N-domain (purple) bound to the VCPIP1 UBX-L (orange). **d**, Overlay of the VCP N-VCPIP1 UBX-L model with an x-ray crystal structure of VCP N bound to OTU1 UBX-L (PDB: 4KDL). VCP N domains were aligned. **e**, Overlay of the VCPIP1 UBX-L model aligned with AlphaFold models for UBP19 (AF-O94966-F1 aa 679-762) and USP9X (AF-Q93008-F1 aa 884-967). **f**, *Top*, Domain map of VCPIP1 ΔUBX-L (aa 1-743). Dashed line indicates residues removed (aa 744-1222). *Bottom,* mass photometry of VCP WT mixed with VCPIP1 ΔUBX-L. **g**, SDS-PAGE following immobilized metal affinity chromatography (IMAC) of His-tagged VCP and different VCPIP1 constructs. Left lanes show input, right lanes show eluate from His dynabeads. VCPIP1 ΔUBX-L is marked with an asterisk. **h**, As in **g** but with varying concentrations of VCPIP1 ΔUBX-L.

Focusing on high resolution regions of the map, we generated a model for the VCP-VCPIP1 asymmetric unit starting from individual domains of a published ADP-bound VCP structure (PDB: 5FTL) (40) and VCP C-terminal tail residues 764 to 775, which were resolved in a peptide substrate-bound VCP structure (PDB: 7LN6) (41). Our map had sufficient resolution to assign and build side chains into the density corresponding to the D1 and D2 domains (Fig. 3b and Supplementary Data Fig. 4c). We observe density for ADP in the D1 domain of both VCP protomers (Supplementary Data Fig. 4e). We do not observe density for nucleotide in VCP D2 domains, suggesting that the D2 domain, which is the main source of VCP’s ATPase activity (42), hydrolyzed and released any bound ATP. Density corresponding to the VCP N-domain is weak, consistent with conformational flexibility, and therefore we did not include the N-domain in this model at this stage (see below).

To model VCPIP1, we started from an AlphaFold prediction (AF-Q96JH7-F1) (43, 44) (see Methods). We identify a VCPIP1 “stalk” near the exit of the VCP central pore (Fig. 3a). Density for side chains in the stalk region enabled us to build a model for the interaction between the VCP D2 domain and VCPIP1 (Fig. 3b) (see Fig. 4). The density connected to the VCPIP1 stalk domain, which we predict is the OTU domain, is highly flexible and low resolution and was omitted from this model (see Fig. 4).

**Fig. 4:**
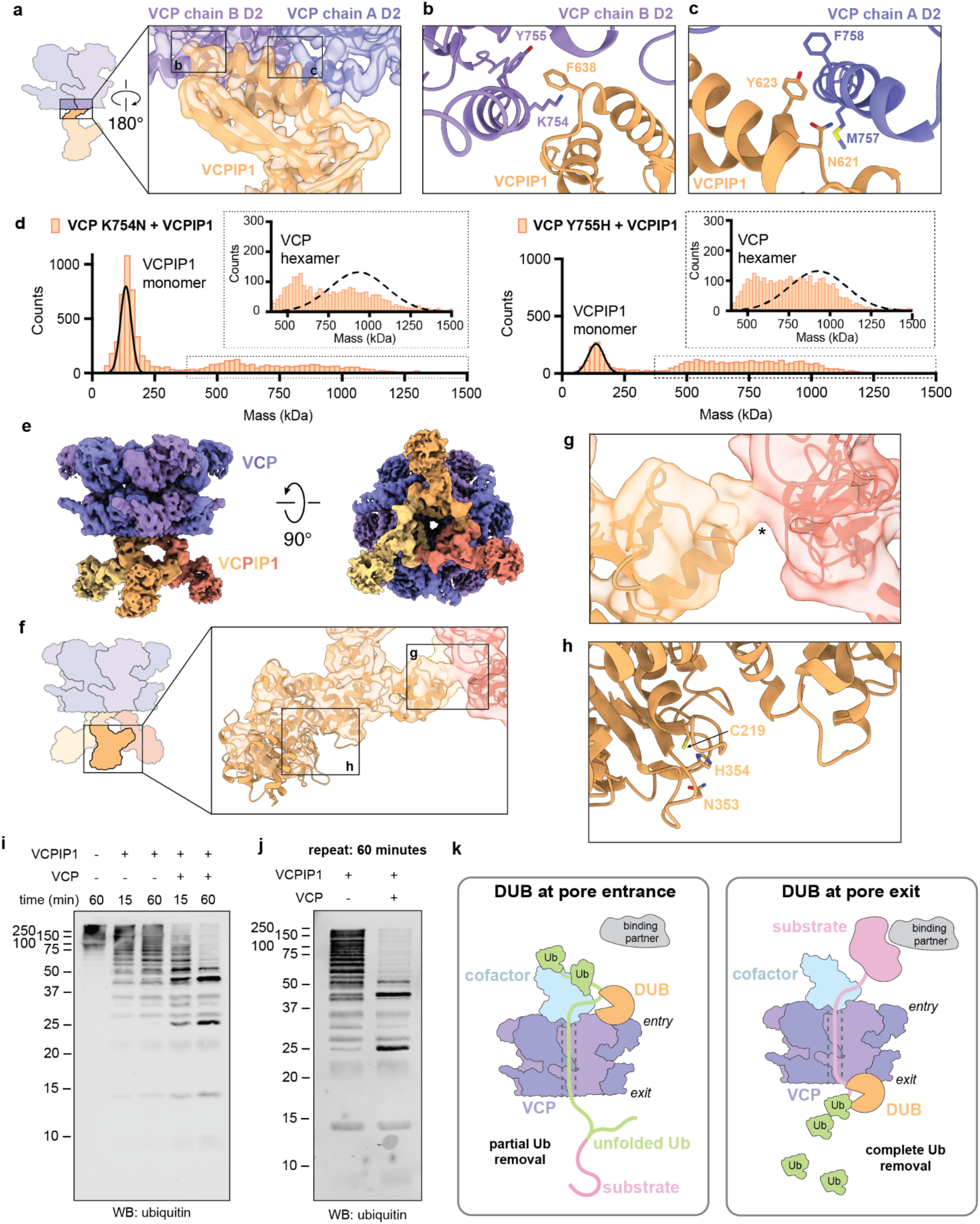
The VCPIP1 deubiquitinase domain is poised to cleave ubiquitin from substrates as they exit VCP’s central pore, while other DUBs can bind at the substrate entry site. **a**, *Left*, Schematic for the asymmetric unit with a box around the VCP D2-VCPIP1 stalk interaction. *Right*, Cryo-EM map and model of two VCP D2 domains (blue and purple) bound to the VCPIP1 stalk (orange). **b**, Zoom of the VCP A D2 (purple)-VCPIP1 stalk (orange) interaction. Key interacting residues are shown in sticks. **c**, as in **b** but the VCP B D2 (blue)-VCPIP1 stalk (orange) interaction. **d**, Mass photometry of mutant VCP-VCPIP1 complexes. *Left*, VCP K754N mixed with VCPIP1 WT. *Right*, VCP Y755H mixed with VCPIP1 WT. The insets (dotted lines) are zooms between masses of 400 and 1500 kDa. The Gaussian distribution (dashed lines) is from VCP WT mixed with VCPIP1 WT (from Fig. 2b). **e**, Side and bottom views of the cryo-EM map of the VCP hexamer (blue and purple) bound to three VCPIP1 protomers (red, orange, yellow). **f**, *Left*, Schematic for the VCP-VCPIP1 complex with a box around the VCPIP1 OTU domain. *Right*, Zoom of the cryo-EM map and model for the VCP hexamer bound to three VCPIP1 protomers. The VCPIP1 OTU domain (orange) is shown. The insets are further zooms of the OTU domain shown in **g** and **h**. **g**, Zoom of the cryo-EM map and model for the VCP hexamer bound to three VCPIP1 protomers. The VCPIP1 OTU domains (orange and red) are shown with connecting density (asterisk). **h**, Zoom of the model for the VCP hexamer bound to three VCPIP1 protomers. The catalytic residues of the VCPIP1 OTU domain (orange) are shown in sticks. **i-j**, Western blots for ubiquitin to measure deubiquitinase activity of VCPIP1 in the presence and absence of VCPIP1. The increase in short ubiquitin chains after 60 minutes was analyzed. VCP stimulates VCPIP1 deubiquitinase activity between 10- and 50-fold across two independent experiments. **k**, Model for coupling of VCP unfoldase and VCPIP1 deubiquitinase activities. VCP (purple) and a cofactor (light blue) recruit a polyubiquitylated substrate (substrate pink, Ub green) along with another binding partner (grey). VCP unfolds ubiquitin and substrate through its central pore (dashed lines). DUBs (orange) bind VCP at either the central pore’s entry or exit sites to remove ubiquitin from substrates. Complete removal of ubiquitin would lead to substrate recycling and partial removal would promote substrate degradation by the proteasome.

Our overall map does not reveal density that can be assigned to VCPIP1 near the entry to VCP’s central pore. Therefore, we performed additional processing of the VCP N-domain (Supplementary Data Fig. 3a). In classes where the N-domain was coaxial with the D1 (‘up’), density only for the VCP N-domain was observed and rigid-body fitted with a model (PDB: 5FTN) (40). When the N-domain was coplanar with the D1 domain (‘down’), we rigid-body fitted VCP with a model (PDB: 5FTL)(40) and observed an additional density (Supplementary Data Fig. 5a). We further refined this class and resolved the structure to ∼4.4 Å resolution (Fig. 3c and Supplementary Data Fig. 5b,c). The additional density was rigid body fit with a model for the VCPIP1 UBX-L domain (AF-Q96JH7-F1) (43, 44) (Fig. 3c). Gratifyingly, our model is consistent with an X-ray crystal structure of the VCP N-domain in complex with the UBX-L of OTU1, a yeast deubiquitinase (PDB: 4KDL), where in both cases the UBX-L VCP-interacting motif (aa GFPP) is positioned in a cleft between N subdomains (31) (Fig. 3d and Supplementary Data Fig. 5d). AlphaFold predictions for DUBs identified in our mass spectrometry datasets (UBP19: AF-O94966-F1, USP9X: AF-Q93008-F1) (43, 44) reveal secondary structures with a similar fold to the VCPIP1 UBX-L (Fig. 3e).

To examine if the VCP N-VCPIP1 UBX-L interaction is required for complex formation, we used MP and immobilized metal affinity chromatography (IMAC). A VCPIP1 mutant lacking its C-terminal UBX-L domain (VCPIP1 ΔUBX-L, aa 1-743) was expressed and purified using the same protocol as wild-type (Fig. 3f and Supplementary Data Fig. 2a). By MP, VCPIP1 ΔUBX-L is monomeric (mean ± SD: 83 ± 14 kDa) (Supplementary Data Fig. 5e). When wild-type VCP was mixed with VCPIP1 ΔUBX-L, we did not observe binding under the conditions of MP experiments but identified gaussian peaks for the proteins alone (mean ± SD: VCPIP1 ΔUBX-L = 83 ± 19 kDa, VCP = 534 ± 69 kDa) (Fig. 3f).

For IMAC analyses, we purified VCP with a hexahistidine tag at its N-terminus (hereafter His-VCP) (Supplementary Data Fig. 2a). We observe nearly all of the input VCPIP1 WT bound to His-VCP-immobilized beads, whereas VCPIP1 ΔUBX-L exhibits reduced binding (Fig. 3g). In an experiment where VCPIP1 ΔUBX-L was titrated (0-1.2 µM), we observe a concentration-dependent increase in VCPIP1 ΔUBX-L bound to His-VCP-immobilized beads (Fig. 3h). Together, these data suggest that the VCPIP1 UBX-L contributes to complex formation by binding VCP in its ADP-bound state, where the N domains adopt a ‘down’ conformation.

### The VCPIP1 deubiquitinase domain is poised to cleave ubiquitin from unfolded substrates at the exit of VCP’s central pore

We next used our map and model of the asymmetric unit (Fig. 3a,b) to identify key interactions between VCPIP1 and the exit of VCP’s central pore. Consistent with up to three VCPIP1 protomers binding the VCP hexamer, our model reveals one VCPIP1 interacting with two VCP protomers (Fig. 4a-c). First, we observe VCPIP1 F638 interacting with K754 and Y755 in the C-terminal helix on one VCP protomer (Fig. 4b). In the adjacent VCP protomer, we observe an interaction interface between VCPIP1 N621/Y623 and VCP C-terminal helix residues M757/F758 (Fig. 4c). Both of these interactions are distinct from VCP interactions with PUB or PUL domains (36).

To determine the contribution of these interactions for complex formation, we purified two VCP mutants, K754N and Y755H, which lack residues that our structural data indicate are important for binding VCPIP1 (Supplementary Data Fig. 2a). Using MP, we observe these mutants as hexamers (Supplementary Data Fig. 6a). When VCP mutants were mixed with wild-type VCPIP1, we observed peaks for VCPIP1 monomer and VCP hexamer (Fig. 4d). More unbound VCP hexamer is present in comparison to wild-type VCP (Figs. 2b and 4d). We also observe broad, low intensity peaks for VCP-VCPIP1 complexes (Fig. 4d). These data suggest that the VCP mutants exhibit weaker binding to VCPIP1.

To examine the flexible region of VCPIP1 that connects to its stalk, we processed a map containing three VCPIP1 protomers bound to the VCP hexamer (Fig. 2d). Again, we performed C3 symmetry expansion and local refinement to obtain a ∼3.4 Å resolution map (Fig. 4e and Supplementary Data Figs. 3a and 6b,c). The OTU domain is estimated to be ∼5 Å resolution, which was sufficient to rigid body fit an AlphaFold model for VCPIP1 (AF-Q96JH7-F1, aa 112-553) (43, 44) into the density (Fig. 4f and Supplementary Data Fig. 6c). The overall shape of the model corresponds well with the density and we are able to identify secondary structural elements (Fig. 4f). We identified two key characteristics of the OTU domain. First, we observe density connecting the OTU domains of adjacent VCPIP1 protomers, but are unable to assign this at the current resolution (Fig. 4f,g). Second, we observe the deubiquitinase active site, consisting of the catalytic triad of C219, H354, and N353, exposed to a cleft where we suspect ubiquitin may bind (Fig. 4f,h). Our cryo-EM map containing three VCPIP1 protomers suggests that multiple deubiquitinase domains can be positioned at the exit of the VCP central pore and their activity may be stimulated through interdomain interactions.

We next tested VCPIP1 DUB activity in the presence of VCP. We performed a Western blot-based experiment using tagless WT VCPIP1 and VCP, and a model substrate, K48-linked poly-ubiquitinated and photocleaved mEos3.2 (hereafter, polyUb_2_-mEos) (45) (Fig. 4i and Supplementary Data Fig. 6d,e). In the absence of VCPIP1, we observe unprocessed polyUb_2_-mEos over the course of the experiment, where several ubiquitin molecules are covalently linked to mEos (Fig. 4i). Addition of VCPIP1 resulted in the reduction of higher molecular weight bands and the appearance of lower molecular weight bands over time, consistent with the loss of ubiquitin chains (Fig. 4i,j). In the presence of VCP, we observe even further reduction in longer mEos-linked ubiquitin chains and increased presence of shorter chains (Fig. 4i,j). As this experiment was performed in the absence of ATP or cofactors, stimulation of VCPIP1 DUB activity by VCP is ATPase and unfoldase independent. Together, these data reveal an interaction at the exit of the VCP central pore that facilitates DUB activation and positions the VCPIP1 catalytic domain for ubiquitin cleavage as substrates exit the pore.

## Discussion

Our proteomics workflow identifies proteins that directly bind at the entry or exit sites of VCP’s central pore in living human cells, and find that the DUB VCPIP1 binds at both sites. We also report the first structural model for how the active sites in VCP and a DUB are coupled. The VCPIP1 UBX-L domain binds the VCP N-domain and the VCPIP1 “stalk” binds proximal to the VCP C-terminal tail, an interaction distinct from other regulatory VCP cofactors (36). This conformation positions the VCPIP1’s deubiquitinase domain below the exit of VCP’s central pore, through which unfolded substrates are likely to emerge. Additionally, binding of VCPIP1 to VCP stimulates its DUB activity against K48-linked ubiquitylated substrates.

Based on our findings, we propose a model for how VCP’s unfoldase activity can be differentially coupled to distinct DUBs (Fig. 4k). Previous studies, many of which focus on the VCP ortholog Cdc48, indicate the unfolding or extraction of substrates is initiated through the recognition of 3 or more ubiquitins by the VCP/UN complex, followed by insertion of one ubiquitin into VCP’s central pore (8, 32). VCP uses ATP hydrolysis to unfold ubiquitin molecules positioned proximal to the substrate and the substrate itself. DUBs at the entry site recognize folded ubiquitins, thus cleaving those more distal to the substrate that is being unfolded. Translocated, polyubiquitin-linked substrates need to remain unfolded to be degraded, which may occur through interactions with chaperones or direct transfer to the proteasome by shuttling factors (Fig. 4k, left) (25). While additional structural information for VCP-DUB interactions is needed, our AlphaFold modeling suggests most DUBs that bind VCP exhibit well-characterized interactions with the VCP N-domain (Fig. 3e). In the case of VCPIP1, up to three protease sites are positioned at the exit of the pore and our analyses do not suggest that this interaction is mimicked by other DUBs (Fig. 4k, right). After translocation of polyubiquitylated substrates through VCP’s central pore, ubiquitins linked to the unfolded substrate must refold before recognition and cleavage by VCPIP1, whose activity is stimulated by binding to VCP. This would require a delay, which could be achieved through the VCPIP1 UBX-L domain binding to VCP’s N-domain and stabilizing the ADP-bound state. Complete removal of ubiquitin from substrate would allow recycling of the proteins removed from multiprotein complexes or membranes. In particular, this mechanism may be utilized in post-mitotic Golgi and ER membrane biogenesis, in which it is proposed that SNARE complexes are recycled by a VCP/VCPIP1/p47 complex (11, 46, 47).

Our structural data show that the VCP-VCPIP1 unfoldase-deubiquitinase complex mimics other ATP-dependent proteolysis systems such as the ClpXP and LonP complexes and the proteasome, which also have protease sites positioned where substrates emerge from ATPase mechanoenzymes (48–50). Unlike these systems, the different VCP-DUB binding modes revealed by our proteomics data would allow polyubiquitylated substrates to be recycled or degraded. Ubiquitylation serves as an essential signal for many biological pathways, including proteostasis (i.e., degradation and autophagy), cell signaling (e.g., NF-kB) and cell division, and the complexity of ubiquitin modifications, such as linkage type and length, enables robust regulation of these dynamic processes (1, 26). Further, the accumulation of misfolded proteins or aggregates is linked to aging and neurodegenerative diseases, and cancer cells can be uniquely susceptible to inhibition or dysfunction of the ubiquitin proteasome system (51), underscoring the importance of faithful ubiquitin-dependent substrate protein processing for maintenance of cellular homeostasis. Our data suggest that VCP-DUB coupling provides important tunable regulation of distinct outcomes for ubiquitylated substrates that require force-dependent extraction or unfolding prior to additional processing.

## Methods

### Plasmids and molecular cloning

Amber suppression plasmid pCMV-AbK was a gift from Peter Schultz (Scripps) (52). Mammalian expression plasmids encoding amber-containing VCP with tandem C-terminal 12X-His and 3X-FLAG tags were generated by Gibson assembly (53) into the pCDNA3.1 vector (ThermoFisher cat. No. V80020). Plasmid for expressing wild-type human VCP in bacteria was obtained from T.-F. Chou (Caltech). To express VCPIP1 in bacteria, the wild-type human VCPIP1 gene was purchased from Addgene (plasmid #22592 from Wade Harper) (28), PCR amplified, and cloned into a pET15b vector using Gibson assembly. Plasmids for VCP and VCPIP1 point mutants were generated by site-directed mutagenesis. VCPIP1 ΔUBX-L (aa 1-743) was generated by PCR amplification from the VCPIP1 full-length gene and Gibson assembly into a pET15b vector. Plasmids for expression of human His-tagged Ufd1 and tagless Npl4 were purchased from Addgene (plasmid #117107 (Ufd1-His) and #117108 (Npl4) from Hemmo Meyer) (33). Plasmids for the synthesis and purification and of polyUb_2_-mEos were purchased from Addgene (ubiquitin: plasmid #12647 from Rachel Klevit (54); mouse Ube1: plasmid #32534 from Jorge Eduardo Azevedo (55)) or a gift from Andreas Martin (plasmids encoding genes for His_6_-Ub_G76V_-Ub_G76V_-mEos3.2 and a gp78-Ubc7 fusion protein) (45). A plasmid encoding only one ubiquitin linearly fused to mEos3.2 (His_6_-Ub_G76V_-mEos3.2) was generated by PCR amplification from this plasmid and Gibson assembly into a pET28a vector.

### Amino acids

3’azibutyl-*N*-carbamoyl-lysine (AbK), was synthesized as described previously (52). *N^ε^*-Boc-lysine (BocK) was purchased from Chem-Impex International (cat. No. 00363).

### Cell culture

HEK293T cells were grown at 37 °C in a humidified atmosphere with 5% CO_2_ in Dulbecco’s Modified Eagle Medium (DMEM) (ThermoFisher cat. No. 11995065) containing 10% fetal bovine serum (Sigma Aldrich cat. No. F4135).

### Transfections/photo-crosslinking

Cells were transfected with the plasmid for VCP(TAG)-12xHis-3xFLAG and pCMV-AbK using lipofectamine 2000 (ThermoFisher cat. No. 11668027) according to manufacturer’s instructions. After an overnight incubation with transfection reagent, the medium was replaced and supplemented with 0.5 mM AbK or 1 mM BocK. 24 hours later, the medium was replaced with cold 1x DPBS (ThermoFisher cat. No. 14190144) and cells were exposed to 365 nm UV light (Spectroline ML-3500S) for 15 minutes on ice. Cells were harvested in lysis buffer (His pulldown only: 20 mM HEPES pH 7.5, 150 mM KCl, 20 mM imidazole, 10 mM beta-mercaptoethanol (βME), 0.2% Triton X-100, 1 mM phenylmethylsulfonyl fluoride (PMSF), and cOmplete EDTA-free protease inhibitor cocktail (Sigma cat. No. 11873580001); tandem immunoprecipitation: 20 mM HEPES pH 7.5, 150 mM KCl, 10 mM βME, 0.2% Tween-20, 1 mM PMSF, and cOmplete EDTA-free protease inhibitor cocktail) and incubated on ice for 15 minutes. The suspension was centrifuged at 16,100g for 15 minutes at 4 °C and the supernatant was collected, flash frozen in liquid nitrogen, and stored at -80 °C until affinity purification.

### Preparation of soluble extracts and enrichment of photo-crosslinked proteins

To prepare soluble extracts using metal affinity purification only, cell lysates were thawed, diluted 2-fold with 8 M urea, and incubated with His-tag Isolation and Pulldown Dynabeads (ThermoFisher cat. No. 10103D) at room temperature for 1 hour. Beads were washed three times with 20 mM HEPES pH 7.5, 1 M KCl, 20 mM imidazole, 10 mM βME, 0.1% Triton X-100, and 8 M urea. Proteins were eluted in 20 mM HEPES pH 7.5, 150 mM KCl, 500 mM imidazole, 10 mM βME, 0.1 % Triton X-100 and 8 M urea at 70 °C for 10 minutes with agitation. For preparation of soluble extracts using tandem immunoprecipitation, cell lysates were thawed and incubated with Pierce Anti-DYKDDDK Magnetic Agarose (ThermoFisher cat. No. A36797) at room temperature for 30 minutes. The agarose was washed three times with 20 mM HEPES pH 7.5, 500 mM KCl, 10 mM βME, 0.2% Tween-20. Proteins were eluted in 20 mM HEPES pH 7.5, 150 mM KCl, 20 mM imidazole, 10 mM βME, 0.2% Tween-20, and 8 M urea at 37 °C for 10 minutes with agitation. The eluate was then incubated with His-tag Isolation and Pulldown Dynabeads at room temperature for 30 minutes. Beads were washed with 20 mM HEPES pH 7.5, 1 M KCl, 20 mM imidazole, 10 mM βME, 0.1% Tween-20, and 8 M urea. Proteins were eluted in 20 mM HEPES pH 7.5, 150 mM KCl, 500 mM imidazole, 10 mM βME, 0.1 % Tween-20 and 8 M urea at 70 °C for 10 minutes with agitation. Eluates were either used directly for Western blot analysis or flash frozen in liquid nitrogen and stored at -80 °C until mass spectrometry sample preparation.

### Antibodies

For Western blot analysis, the following primary antibodies were used: anti-FLAG (Sigma cat. No. F1804), anti-VCP (Santa Cruz Biotechnology cat. No. sc-57492), anti-Ub (Santa Cruz Biotechnology cat. No. sc-8017), and anti-GAPDH (Proteintech cat. No. 10494-1-AP). IRDye-conjugated secondary antibodies raised in goat or donkey were purchased from LI-COR Biosciences (cat. Nos. 925-68070 and 925-32213) and used according to the manufacturer’s instructions.

### Mass spectrometry

Protein samples were mixed with NuPAGE LDS sample buffer (ThermoFisher cat. No. NP0007) and reduced with 50 mM DTT at 70 °C for 10 minutes with agitation. Samples were cooled to room temperature, alkylated with 100 mM iodoacetamide for 30 minutes, and electrophoresed on a NuPAGE 10% Bis-Tris gel (ThermoFisher cat. No. NP0302BOX) only until the entire sample entered the gel (∼3-4 minutes at 130 V). The gel was stained with GelCode Blue reagent (ThermoFisher cat. No. 24590) and destained with water. Proteins in gel plugs were digested, and peptides were extracted and purified as described (56). Tryptic peptides were analyzed by LC-MS using an Orbitrap Exploris mass spectrometer coupled with an Easy-nLC system (Thermo Fisher Scientific). SpectroMine (Biognosys AG) software was used for label-free quantification (LFQ) analyses and the protein LFQ outputs from SpectroMine were further analyzed using Microsoft Excel. To compare the LFQs between AbK- and BocK-treated samples, normalization was applied so that the LFQs for VCP were equal.

### Protein purification

Full-length human tagless and His-tagged VCP were expressed and purified as described (18), but the TEV protease cleavage step was omitted from the purification of His-tagged VCP.

Full length human VCPIP1 (WT and variants) was expressed and purified as described (11), but with modifications. VCPIP1 was expressed in *E. coli* Rosetta (DE3) pLysS cells (Merck, cat. No. 70954) and grown following the same procedure used for VCP. Cells were pelleted and resuspended in lysis buffer (50 mM Tris-HCl pH 7.4, 300 mM KCl, 20 mM imidazole, 5% glycerol, 2 mM β-mercaptoethanol, 1 mM *p*-phenylmethylsulfonyl fluoride, cOmplete EDTA-free protease inhibitor cocktail (Roche)). All subsequent purification steps were performed at 4 °C. Cells were lysed using an EmulsiFlex C5 homogenizer (Avestin, ∼6 cycles at 10,000-15,000 psi homogenization pressure). The lysate was clarified by centrifugation at 45,000 rpm for 30 min using a Type 70 Ti rotor in a Beckman Coulter Optima LE-80K ultracentrifuge. The clarified lysate was filtered and loaded onto Ni-NTA resin and incubated for 1 hr. The resin was washed with ∼200 mL of wash buffer (50 mM Tris-HCl pH 7.4, 300 mM KCl, 20 mM imidazole, 5% glycerol, 2 mM βME) and eluted with elution buffer (50 mM Tris-HCl pH 7.4, 300 mM KCl, 300 mM imidazole, 5% glycerol, 2 mM βME). The eluate was treated with tobacco etch virus (TEV) protease (∼1 mg) and dialyzed in dialysis buffer (50 mM HEPES pH 7.5, 150 mM KCl, 20 mM imidazole, 5% glycerol, 2 mM βME) overnight. Protein was passed 3 times over Ni-NTA resin pre-equilibrated with reverse Ni buffer (50 mM HEPES pH 7.4, 150 mM KCl, 20 mM imidazole, 5% glycerol, 2 mM βME), or incubated 1 hr then eluted. The flow through was concentrated using an Amicon Ultra 100K concentrator to 0.5 mL, centrifuged, and loaded into a Superdex 200 Increase 10/300 column (GE Healthcare) equilibrated with gel filtration buffer (50 mM HEPES pH 7.5, 150 mM KCl, 1 mM DTT). The fractions containing purified VCPIP1 were pooled, concentrated using an Amicon Ultra 100K concentrator, and frozen in liquid nitrogen and stored at -80 °C.

Purifications of mono-Ub, Ube1, gp78-Ubc7, and His_6_-Ub_G76V_-Ub_G76V_-mEos3.2 were performed as described previously (45, 57). To purify His_6_-Ub_G76V_-mEos3.2, the protocol for purification of His_6_-Ub_G76V_-Ub_G76V_-mEos3.2 was followed. To synthesize and purify polyUb_1_-mEos or polyUb_2_-mEos, His_6_-Ub_G76V_-mEos3.2 or His_6_-Ub_G76V_-Ub_G76V_-mEos3.2 (20 µM) was mixed with Ube1 (1 µM), gp78-Ubc7 (10 µM), and ubiquitin (400 µM) in 2 mL buffer (20 mM HEPES pH 7.4, 150 mM KCl, 5 mM MgCl_2_, 10 mM MgATP). Ubiquitin was added sequentially every hour for the first 6 hours of incubation at 37 °C. After the final addition of ubiquitin, the reaction was kept at 37 °C overnight (total = 23 hours). The following day, the sample was placed on ice and incubated under UV light (365 nm) for 90 minutes for photo-conversion of mEos. The ubiquitinated, photo-converted substrate was incubated with Ni-NTA resin for 2 hours at 4 °C. The resin was washed with 50 mL of wash buffer (20 mM HEPES pH 7.4, 250 mM KCl, 1 mM MgCl_2_, 20 mM imidazole, 5% glycerol, 2 mM β-mercaptoethanol) and eluted with elution buffer (20 mM HEPES pH 7.4, 250 mM KCl, 1 mM MgCl_2_, 300 mM imidazole, 5% glycerol, 2 mM β-mercaptoethanol). The elution was concentrated using an Amicon Ultra 10K concentrator to 0.5 mL, centrifuged, and loaded into a Superose 6 Increase 10/300 column (GE Healthcare) equilibrated with gel filtration buffer (20 mM HEPES pH 7.4, 250 mM KCl, 1 mM MgCl_2_, 5% glycerol, 1 mM TCEP). The fractions containing long, medium, and short polyUb-mEos chains were pooled, concentrated using an Amicon Ultra 10K concentrator, and frozen in liquid nitrogen and stored at -80 °C.

### Cryo-EM sample preparation

Full-length WT VCP was buffer exchanged into 50 mM HEPES pH 7.5, 25 mM KCl, 2.5 mM MgCl_2_, 2.5 mM GSH using an Amicon Ultra 100K concentrator, and concentrated to ∼1.3 mg/mL. VCP was mixed with VCPIP1 to a final concentration of 10 µM VCP monomer and 20 µM VCPIP1 in the same buffer as above. 3 µL was applied to a glow-discharged Quantifoil R1.2/1.3 Au 400 mesh grid, blotted for 2s, and plunge-frozen into liquid ethane using a FEI Vitrobot IV operated at 4 °C and 100% humidity.

### Cryo-EM data collection

Cryo-EM data were collected on a Titan Krios microscope (FEI), operating at 300 kV and equipped with a Gatan K3-summit detector in super-resolution mode using SerialEM(58). 7,884 movies were recorded at a nominal magnification of 105,000X, corresponding to a calibrated pixel size of 0.847 Å (super-resolution pixel size of 0.4235 Å/pixel). Each exposure was fractionated across 40 frames with a total electron dose of 44.6 e^-^/Å^2^ and a total exposure time of 1.6 s (dose rate of 20 e^-^/pixel/sec). The defocus values ranged from -0.5 to -2.0 μm.

### Cryo-EM data processing

MotionCor2 was used for dose-weighting and to correct interframe movement for each pixel CTF parameters were estimated using CTF estimation in cryoSPARC (v4.4.1 or v4.5.3) Particles were automatically picked (∼2.8 million) and subjected to 2D classification. A subset of classes was selected as references for ab-initio reconstruction. To clean particles, we used the resulting reconstruction as a reference for heterogeneous refinement. One class (∼780,000 particles) resembling the VCP hexamer was subjected to non-uniform refinement. These particles were subjected to heterogenous refinement with a mask for the density below the VCP hexamer. Separately, we performed a local refinement with C3 symmetry followed by symmetry expansion. This increased the effective size of our dataset (∼2.2 million particles), which was used to perform local refinement and 3D classification in RELION-4.0 (61). 3D classification was performed using a mask surrounding the asymmetric unit consisting of two adjacent VCP protomers and one VCPIP1 protomer. This enabled us to remove particles lacking VCPIP1. We then performed a supervised 3D classification using references with and without the density below VCP. The remaining particles (∼690,000) were subjected to CTF refinement and Bayesian polishing. Finally, we removed duplicates and subsequently performed 3D flexible refinement in cryoSPARC (62) to generate an overall map of the VCP-VCPIP1 complex (Supplemental Figure 4A).

To improve resolution in the VCP N domain, we took symmetry expanded particles after focused 3D classification and performed a second focused 3D classification using a mask surrounding the VCP N and D1 domains. Taking the class with the best density for VCPIP1 on the VCP N domain, we performed a local refinement and removed duplicates. A final local refinement was performed to generate a map of the VCP N-D1 bound to the VCPIP1 UBX-L.

### Cryo-EM model building

VCP (PDB: 5FTL and 7LN6) (40, 41) and VCPIP1 (AF-Q96JH7-F1) (43, 44) models were segmented into individual domains [VCP: D1 and D2, C-terminal tail residues 764-775; VCPIP1: stalk (residues 556-666)]. Each segment was rigid body fitted in UCSF Chimera (version 1.18.0) (63) and flexibly fit using ISOLDE in ChimeraX (version 1.8.0) (64, 65). The docked domains were joined to generate overall VCP (monomer) and VCPIP1 models. Real space refinement in Coot (version 0.9.8.5) (66) with planar peptide and transpeptide restraints was used to adjust side chains for a single chain. The refined coordinates for VCP were replicated and refit into the second protomer. The model was further refined using Phenix (version 1.21.1-5286) (67) real-space refinement and problem areas were fixed manually in Coot.

### Mass photometry

Data were collected using a OneMP mass photometer (Refeyn) calibrated with bovine serum albumin (ThermoFisher, cat. No. 23210), beta amylase (Sigma Aldrich, cat. No. A8781-1VL), and thyroglobulin (Sigma Aldrich, cat. No. T9145-1VL). Focus was adjusted using filtered (0.22 µm) buffer (50 mM HEPES pH 7.5, 25 mM KCl, 2.5 mM MgCl_2_, 2.5 mM GSH), then a 2X solution of protein in filtered buffer was added directly, diluting it 2-fold. Final concentrations of VCP and VCPIP1 were 150 nM (monomer) and 100 nM, respectively. Movies were acquired for 6,000 frames (60 s) using AcquireMP software (version 2022 R1) and default settings. Raw data were converted to frequency distributions using DiscoverMP software (version 2022 R1) or Prism 10 (GraphPad) and a bin size of 10 kDa.

### Pulldown assays

To examine the binding of VCPIP1 variants to VCP, His-VCP (500 nM monomer) was immobilized on His-tag Isolation and Pulldown Dynabeads (ThermoFisher cat. No. 10103D) at room temperature for 30 minutes. Beads were washed three times with buffer (50 mM HEPES pH 7.5, 25 mM KCl, 2.5 mM MgCl_2_, 20 mM imidazole, 10 mM βME, 0.1% Tween-20, and 0.1 mg/mL BSA). Beads were resuspended in buffer with WT or ΔUBX-L VCPIP1 (400 nM) and incubated at room temperature for 30 minutes. Proteins were eluted from beads using buffer supplemented with 500 mM imidazole. The reaction was analyzed by SDS-PAGE and Coomassie staining.

### Deubiquitinase assays

For the Western blot-based DUB assay, VCPIP1 WT (200 nM) was incubated with medium length polyUb_2_-mEos chains (20 nM) and VCP WT (300 nM monomer) in 20 µL of buffer (50 mM HEPES pH 7.5, 25 mM KCl, 2.5 mM MgCl_2_, 1 mM DTT, 0.01% Tween-20) for 15 or 60 minutes at 37 °C. The proteolysis of ubiquitin chains was analyzed by Western blot (anti-Ub antibody: Santa Cruz Biotechnology cat. No. sc-8017)

## Acknowledgements

We thank T.-F. Chou for providing the *H. sapiens* VCP plasmid and A. Martin for providing His_6_-Ub_G76V_-Ub_G76V_-mEos3.2 and gp78-Ubc7 plasmids. We also thank P. Schultz for providing the pCMV-AbK plasmid. T.M.K. (GM130234) and B.T.C. (GM109824) are grateful to the NIH for supporting this research. L.E.V. was supported in part by the NIH T32 GM115327 and GM136640 Chemistry-Biology Interface Training Grant to the Tri-Institutional PhD Program in Chemical biology. We are grateful to M. Ebrahim, J. Sotiris, H. Ng, and the Evelyn Gruss Lipper Cryo-Electron Microscopy Resource Center for cryo-EM support.

