## Supplementary Materials for "Distinct modes of coupling between VCP, an essential unfoldase, and deubiquitinases"

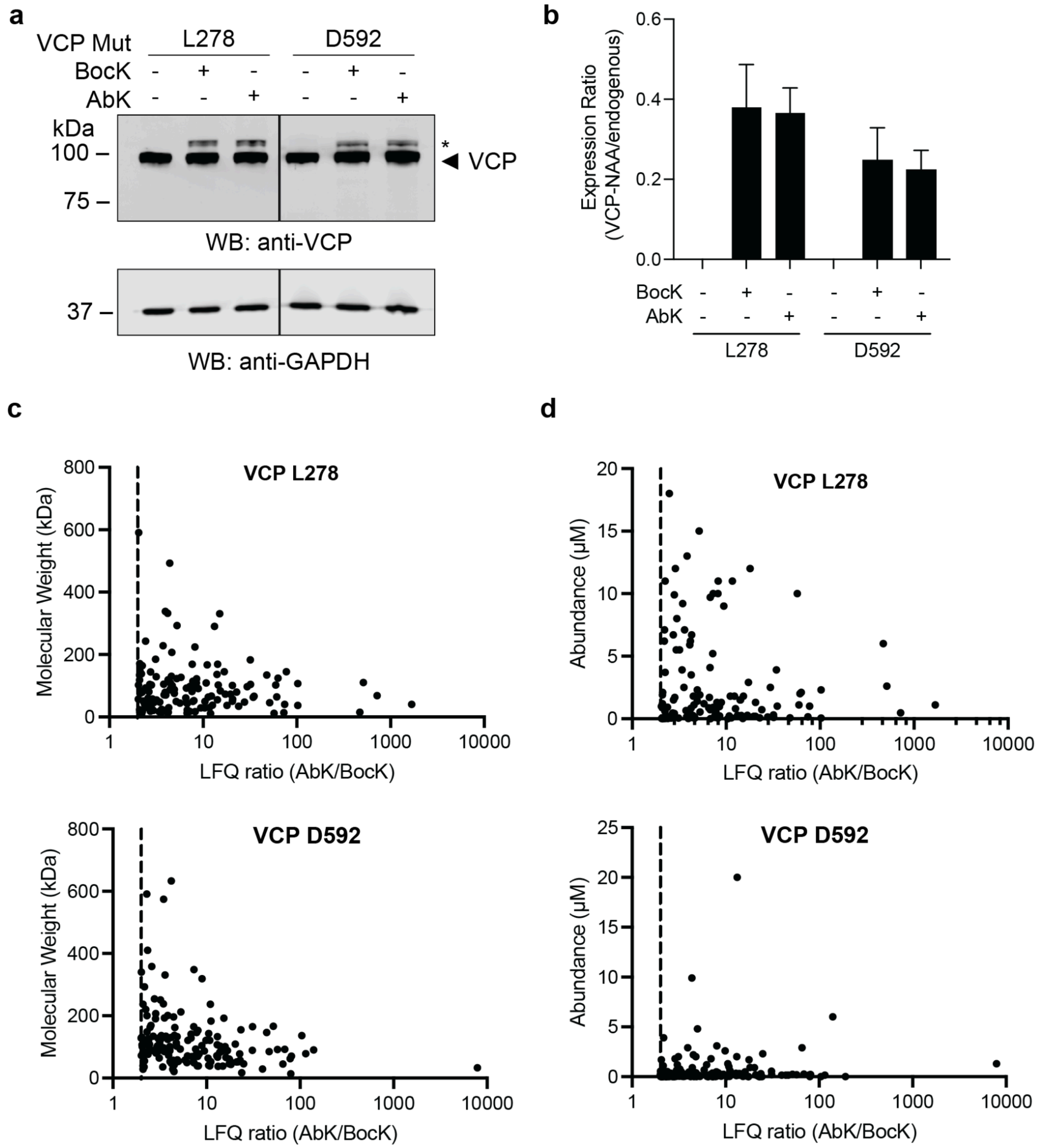


**Supplementary Data Fig. 1: Analysis of the expression of VCP constructs in HEK293T cells and mass spectrometry data.** **a**, Western blot of HEK293T cells expressing VCP with non-natural amino acid (AbK or BocK) incorporated at L278 (VCP-L278-AbK or VCP-L278-BocK) or D592 (VCP-D592-AbK or VCP-D592-BocK) (asterisk, endogenous VCP marked with arrowhead). **b**, Quantification of Western blot in **a**. VCP expressed with a non-natural amino acid (VCP-NAA) is compared to endogenous VCP. (VCP-L278-AbK: 0.365 ± 0.063; VCP-L278-BocK: 0.379 ± 0.108; VCP-D592-AbK: 0.223 ± 0.049; VCP-D592-BocK: 0.247 ± 0.081; mean ± SD; N = 3). **c**, Plots of molecular weight (kDa) versus LFQ ratio (AbK/BocK) for hits (LFQ ratio > 2, dashed line) in mass spectrometry datasets where non-natural amino acids were incorporated at VCP L278 or D592. **d**, Plots of cellular abundance versus LFQ ratio (AbK/BocK) for hits (LFQ ratio > 2, dashed line) in VCP-L278 and VCP-D592 mass spectrometry datasets.

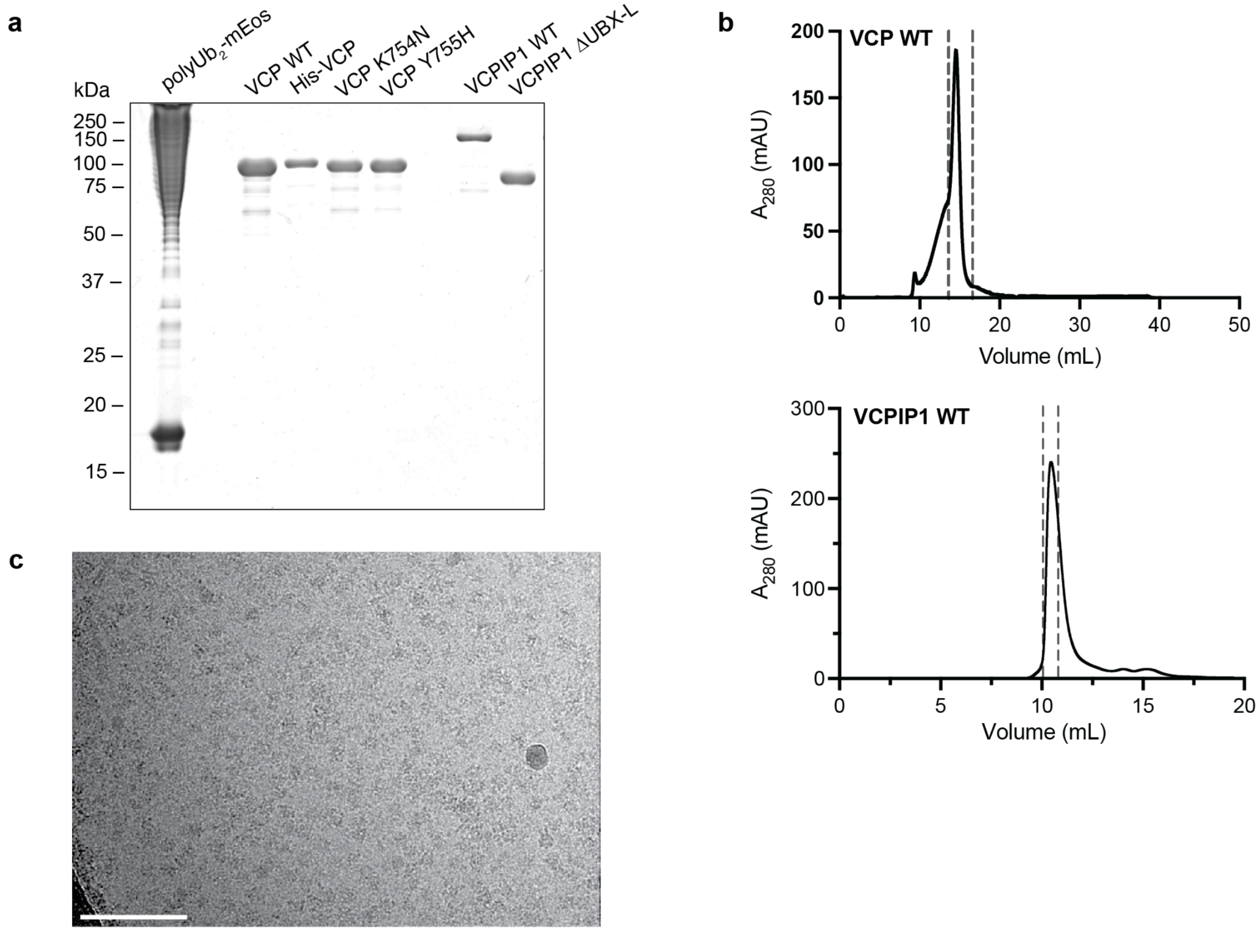


**Supplementary Data Fig. 2: Recombinant protein constructs used in this study and representative cryo-EM micrograph of the VCP-VCPIP1 complex.** **a**, Coomassie-stained SDS-PAGE of protein constructs used in this study. **b**, Representative size exclusion chromatography traces for VCP (top) and VCPIP1 (bottom) WT preps. Dashed lines represent fractions that were collected and pooled. **c**, Example of a cryo-EM micrograph of the VCP-VCPIP1 complex (scale bar = 100 nm).

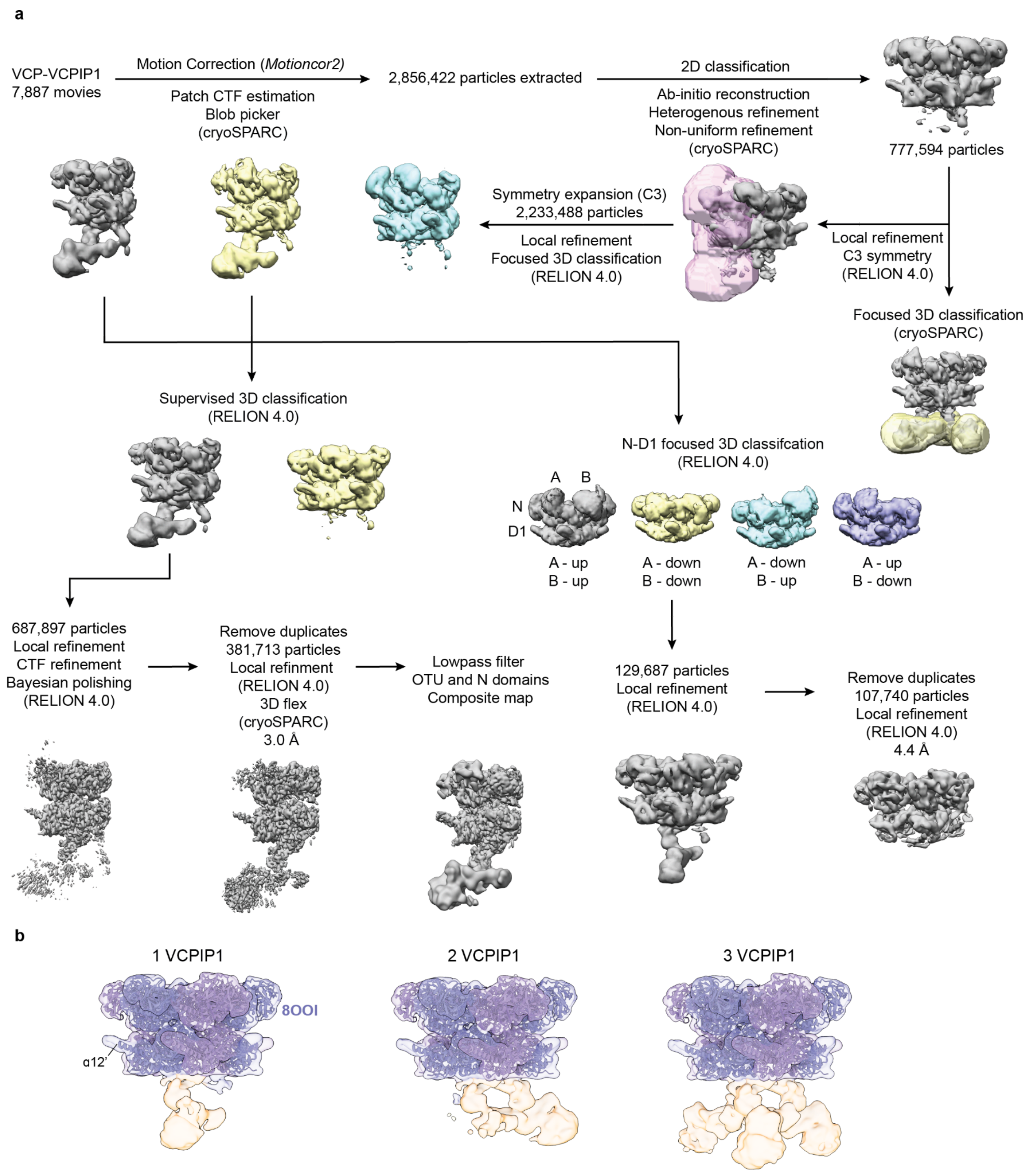


**Supplementary Data Fig. 3: Cryo-EM workflow for the VCP-VCPIP1 complex and initial assignment of density. a**, Cryo-EM workflow for the VCP-VCPIP1 complex. **b**, Side views of 3D classes with 1-3 VCPIP1 protomers (orange) bound to the VCP hexamer (blue and purple) docked with PDB 8OO1. Helix α12' is denoted.

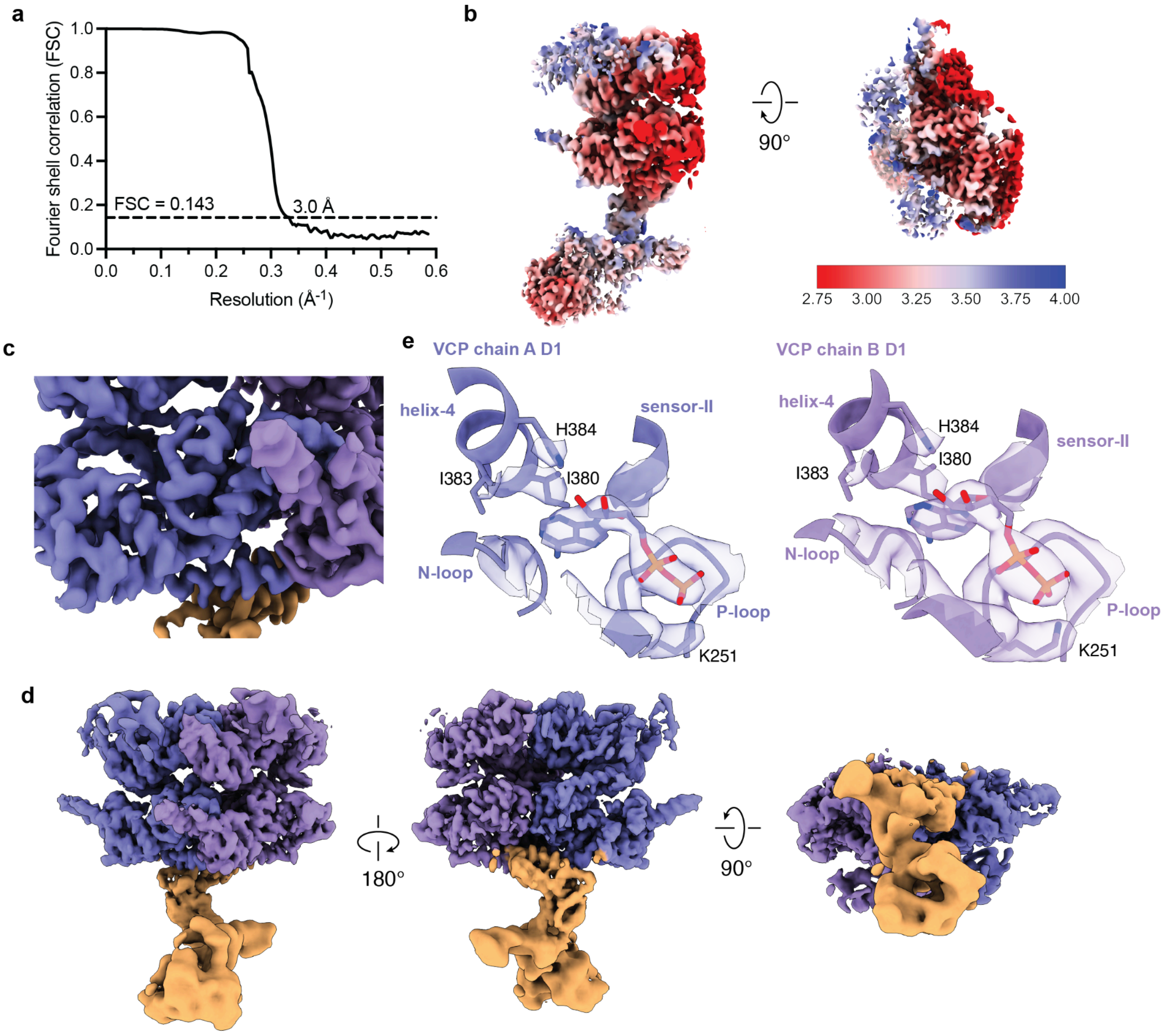


**Supplementary Data Fig. 4: Analysis of the overall VCP-VCPIP1 complex.** **a**, Fourier shell correlation (FSC) curve for the VCP-VCPIP1 complex. **b**, Cryo-EM map of the VCP-VCPIP1 complex colored according to local resolution, showing a distribution from 2.75 to 4.00 Å. **c**, Zoom of cryo-EM map of the VCP (blue and purple)-VCPIP1 (orange) complex showing secondary structure (helices) and side chain resolution. **d**, Zoom of the VCP D1 ATPase domain active sites showing density for ADP. **d**, Composite map of the VCP-VCPIP1 complex in three different views with the VCP N domains and VCPIP1 OTU domain low pass filtered to 7-8 Å resolution.

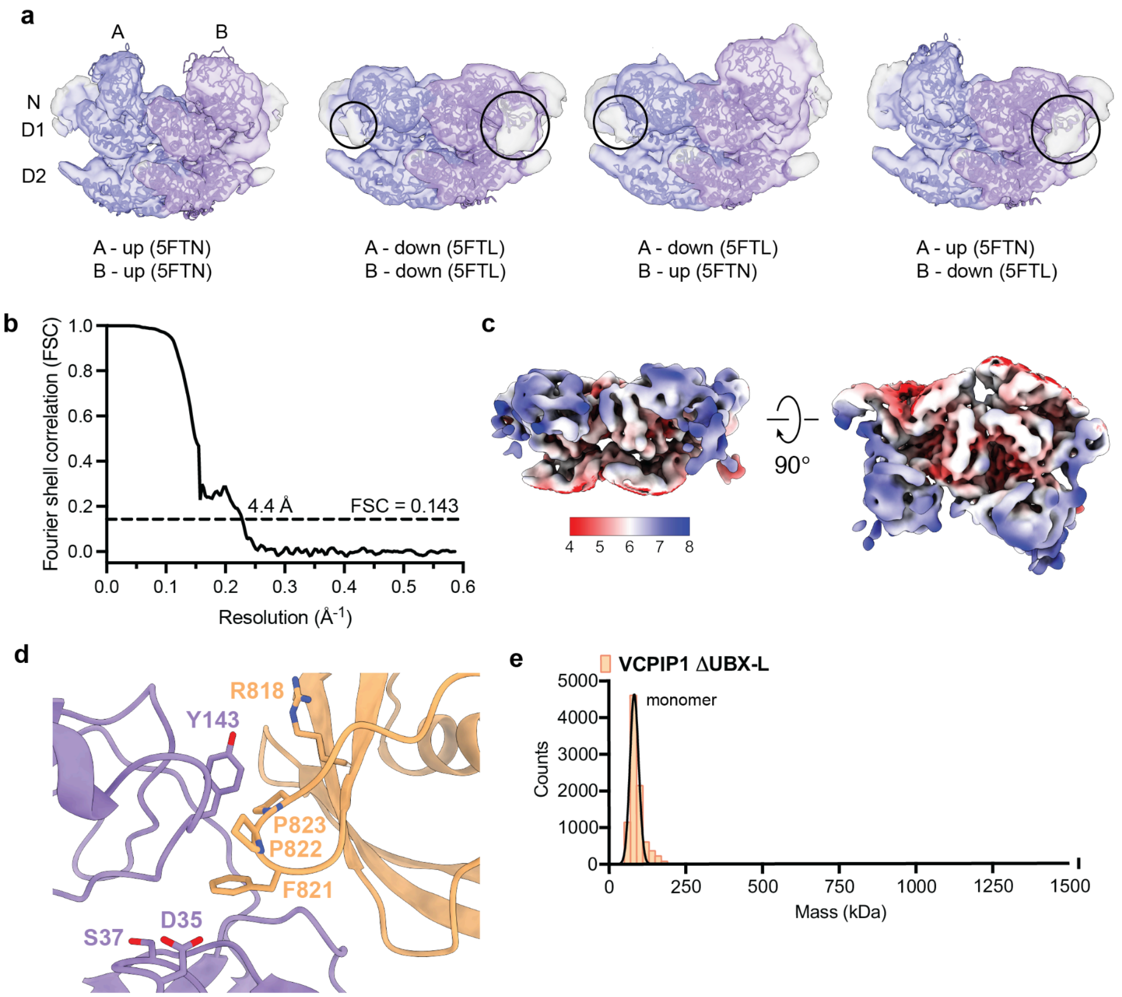


**Supplementary Data Fig. 5: Analysis of the VCP N-VCPIP1 UBX-L interaction. a**, Side views of 3D classes of VCP with the N domain (blue and purple) in 'up' or down' conformations. VCP protomers were rigid body fit with models for VCP with the N domain 'up' (PDB: 5FTN) or 'down' (PDB: 5FTL). An additional density (circled) is observed when N domains are in the 'down' position. **b**, FSC curve for the map of VCP N-VCPIP1 UBX-L. **C**, Cryo-EM map of the VCP N-VCPIP1 UBX-L complex colored according to local resolution, showing a distribution from 4.0 to 8.0 Å. **d**, Zoom of the VCP N-VCPIP1 UBX-L model. Key residues are shown in sticks. **e**, Mass photometry analysis of VCPIP1 ΔUBX-L at 100 nM .

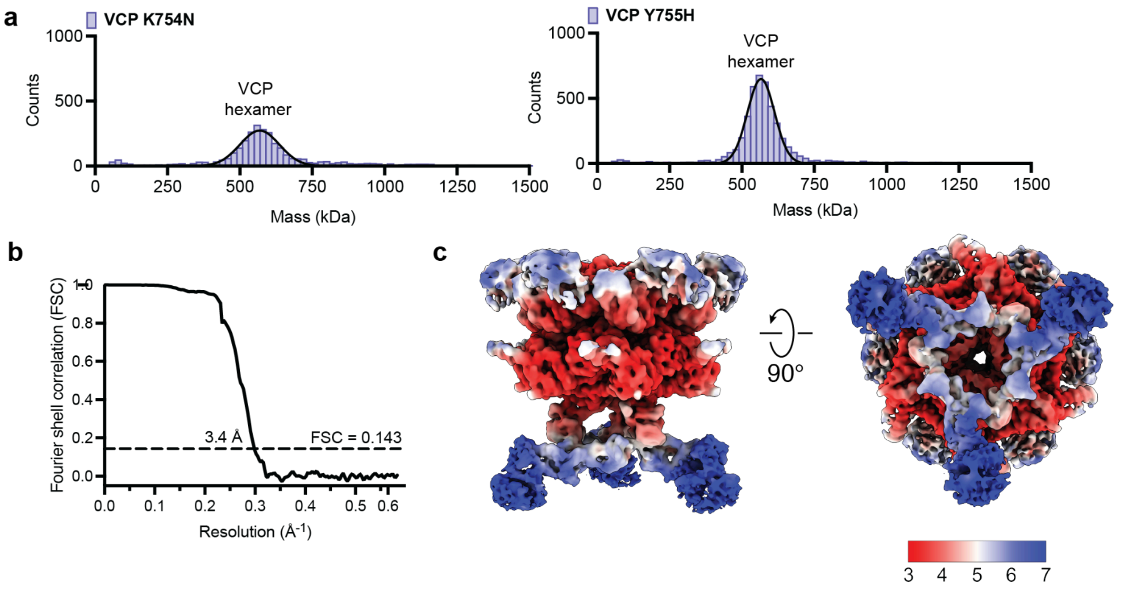


**Supplementary Data Fig. 6: Analysis of the VCP D2-VCPIP1 stalk interaction. a**, Mass photometry analysis of VCP K754N and Y755H mutants. **b**, FSC curve for the VCP hexamer-3 VCPIP1 complex. **c**, Cryo-EM of the VCP hexamer- 3 VCPIP1 complex colored according to local resolution, showing a distribution from 3.0 to 7.0 Å.

**Supplementary Table 1. Mass spectrometry data for selected proteins identified in VCP-L278-AbK and VCP-L278-BocK samples.** LFQ ratios (AbK/BocK), percent coverages, and numbers of unique peptides in run for selected proteins in VCP-L278 samples. See source data for complete list.

| Protein | LFQ ratio (AbK/BocK) | Percent Coverage (AbK, BocK) | Unique Peptides in Run (AbK, BocK) |
|---|---|---|---|
| NSF1C/p47 | 1680.93 | 67.60, 0.00 | 23, 0 |
| NPL4 | 717.17 | 36.50, 0.00 | 19, 0 |
| UBX2B/p37 | 102.46 | 41.40, 0.00 | 13, 0 |
| VCIP/VCPIP1 | 47.62 | 9.70, 0.00 | 11, 0 |
| UFD1 | 23.92 | 7.50, 0.00 | 2, 0 |
| USP9X | 13.22 | 1.90, 0.40 | 4, 1 |
| ALG13 | 0.73 | 4.10, 3.40 | 5, 4 |

**Supplementary Table 2. Mass spectrometry data for selected proteins identified in VCP-D592-AbK and VCP-D592-BocK samples.** LFQ ratios (AbK/BocK), percent coverages, and numbers of unique peptides in run for selected proteins in VCP-L278 samples. See source data for complete list.

| Protein | LFQ ratio (AbK/BocK) | Percent Coverage (AbK, BocK) | Unique Peptides in Run (AbK, BocK) |
|---|---|---|---|
| UBXN1 | 7888.03 | 9.40, 0.00 | 2, 0 |
| UBP19 | 43.89 | 1.50, 0.00 | 2, 0 |
| PLAP/PLAA | 14.77 | 49.80, 1.10 | 34, 1 |
| UBE4B | 7.85 | 7.60, 3.50 | 9, 3 |
| VCIP/VCPIP1 | 3.68 | 29.60, 5.30 | 31, 6 |
| UBP34 | 1.86 | 1.20, 0.30 | 3, 1 |
| UBP24 | 1.44 | 1.10, 0.70 | 3, 2 |
| ALG13 | 1.16 | 6.00, 4.90 | 8, 6 |
| USP9X | 1.02 | 10.60, 7.60 | 26, 18 |
| UBP7 | 0.75 | 0.00, 1.20 | 0, 1 |
| OTU1 | 0.04 | 0.00, 2.60 | 0, 1 |

**Supplementary Table 3. Single-particle cryo-EM data collection, refinement, and validation.** Cryo-EM data collection, refinement, and validation statistics for VCP-VCPIP1.

| | VCP-VCPIP1<br>(EMD-46912, PDB 9DIL) |
|---|---|
| **Data collection and processing** | |
| Microscope | Titan Krios |
| Voltage (kV) | 300 |
| Detector | K3 Summit |
| Magnification | 105,000 |
| Electron exposure ($e^-$/Å$^2$) | 44.6 |
| Exposure rate ($e^-$/pixel/s) | 20 |
| Calibrated pixel size (Å) | 0.847 |
| Defocus range (μm) | -0.5 to -2.0 |
| Particle images (no.) | 7,884 |
| Map resolution (Å) | 3.0 |
| FSC threshold | 0.143 |
| Map sharpening B-factor (Å$^2$) | -73.92 |
| **Refinement** | |
| Model composition | 2 VCP D1/D2, 1 VCPIP1 ‘ stalk’ |
| PDB | 9DIL |
| Non-hydrogen atoms | 9,737 |
| Protein residues | 1,237 |
| Ligands | 2 ADP |
| **R.M.S. deviations** | |
| Bond lengths (Å) | 0.004 |
| Bond angles (°) | 0.492 |
| **Validation** | |
| MolProbity score | 1.69 |
| Clashscore | 4.32 |
| Rotamer outliers (%) | 1.64 |
| **Ramachandran plot** | |
| Favored (%) | 95.68 |
| Allowed (%) | 4.32 |
| Outliers (%) | 0.00 |
